# Test-retest reliable and site-robust Hidden Markov Model framework for discovering whole-brain beta activity

**DOI:** 10.64898/2026.05.07.723415

**Authors:** Sara Korkealaakso, Christine Ahrends, Mia Liljeström, Diego Vidaurre, Hanna Renvall, K Amande M Pauls

**Affiliations:** BioMag Laboratory, HUS Medical Imaging Center, Helsinki University Hospital, Helsinki University and Aalto University School of Science, HUS Helsinki, Finland; Department of Neuroscience and Biomedical Engineering, Aalto University School of Science, Espoo, Finland; Oxford University Centre for Integrative Neuroimaging, Nuffield Department of Clinical Neurosciences, University of Oxford, Oxford, United Kingdom; Center of Functionally Integrative Neuroscience, Department of Clinical Medicine, Aarhus University, Aarhus, Denmark; Linacre college, University of Oxford, Oxford, United Kingdom; Aalto NeuroImaging, Aalto University, 00076 Aalto, Finland; Department of Psychiatry, University of Oxford, Oxford, United Kingdom; Centre de Recerca Matematica, Barcelona, Spain; Department of Neurology, Helsinki University Hospital and Department of Clinical Neurosciences (Neurology), University of Helsinki, Helsinki, Finland

## Abstract

Sensorimotor beta activity (13-30 Hz) is a key neuronal signature in the human sensorimotor system, and its features can be effectively measured using functional brain imaging methods such as magnetoencephalography (MEG). In addition to its importance in healthy brain processing, beta activity has been shown to be altered in several neurological diseases, underscoring its potential as a biomarker. To serve as biomarkers, features must be reliably defined, stable across measurements and, ideally, amenable to automated analysis, yet current approaches to beta characterization require subjective decisions and manual work. We here describe a hidden Markov model (HMM) based approach to automatically segment beta events from source level MEG beta band activity into discrete high- and low-beta states. We demonstrate the differences between the proposed HMM based approach and a commonly used amplitude-envelope based approach to analyse high- and low-beta modulation. We show that the methods complement each other both when applied to resting data and task related passive movement data. Furthermore, we assess the test-retest reliability of the proposed pipeline within individuals using intraclass correlation coefficients (ICC), and test if HMM constructed at one measurement site can be applied to data acquired at another site, thereby evaluating its multisite transferability. We show that the proposed approach produces stable results within subjects and across sites for many of the features. The ICC values were excellent for high-beta state (86-100% of brain areas), while low-beta state test-retest reliability was more modest. Most of the features showed statistically significant differences between sites only in a few brain areas, indicating very good multisite stability. The proposed approach can serve as an automated, reproducible analysis pipeline for, e.g., clinical applications, and appears suitable for multi-site datasets.

## 1. Introduction

Magnetoencephalography (MEG) and electroencephalography (EEG) enable non-invasive measurements of human brain oscillations. One of the first brain cortical rhythms to be characterized was beta activity (13-30 Hz) (Berger, 1929; Hari, 1997). Since then, beta has been extensively investigated in the human motor system, as it is particularly prominent in the sensorimotor cortex and modulated by motor tasks (Jurkiewicz et al., 2006; Salmelin & Hari, 1994). More recent research also points towards a more general functional role of beta activity (Schmidt et al., 2019), including in attention (Haegens et al., 2011), working memory, and executive control (Swann et al., 2009). Beta activity is altered in several neurological conditions, such as Parkinson’s disease (Pauls et al., 2022; Vinding et al., 2020), motor stroke (Bartur et al., 2019; Laaksonen et al., 2012; Parkkonen et al., 2018) and amyotrophic lateral sclerosis (ALS) (Dukic et al., 2022; Proudfoot et al., 2017). Thus, beta band activity plays not only a central role in healthy brain processing, but has potential to serve as an indicator of abnormal brain function in clinical applications, also due to its demonstrated reproducibility and reliability (Illman et al., 2022; Niemelä et al., 2025; Pauls et al., 2024; Piitulainen et al., 2018).

Beta oscillatory activity is not continuous, but emerges in transient ‘bursts’ or ‘events’ (Feingold et al., 2015; Little et al., 2019; Shin et al., 2017). Movement onset is associated with a suppression of beta-band power, referred to as movement-related beta desynchronization (MRBD), which is followed by a post-movement increase in beta activity known as post-movement beta rebound (PMBR) (Pfurtscheller & Lopes Da Silva, 1999). These movement-related changes reflect modulations in burst probabilities (Feingold et al., 2015). Beta bursts are frequently defined as temporary increases in beta band amplitude (Little et al., 2019; Pauls et al., 2024; Tinkhauser et al., 2018). While the amplitude-based approach to detecting bursts yields reproducible burst characteristics (Pauls et al., 2024), it typically involves manual effort and depends on subjective, observer-dependent decisions. Amplitude-thresholding based approaches are also susceptible to differences in signal-to-noise ratio. Moreover, determining an individual beta band based on the dominant peak in the power spectral density (PSD) assumes that beta bursts are limited to a specific frequency, precluding the investigation of multiple beta frequencies.

Hidden Markov Models (HMMs) provide a framework for characterizing brain activity by segmenting multidimensional MEG data into discrete oscillatory states (Quinn et al., 2018; Vidaurre et al., 2016a) defined by, e.g., the signals’ spatial, spectral, and amplitude features (Vidaurre, Hunt, et al., 2018). HMMs, and specifically its time-delay embedded version (TDE-HMM), have recently been utilized to identify beta bursts within broad band (1-48 Hz) MEG activity (Seedat et al., 2020). These studies have indicated that automated HMM approaches can effectively locate beta activity maxima to the sensorimotor regions as expected, and, notably, identify mostly the same events as manual amplitude thresholding-based method, suggesting that characterizing beta activity using HMMs may provide both a data-driven and automated alternative to traditional approaches. However, previous studies have considered beta as a single frequency band, even though there is mounting evidence that beta comprises two separate bands, low-beta and high-beta, with different functional roles (Barone & Rossiter, 2021; Cao et al., 2024, Nougaret et al. 2024). Additionally, previous analyses have been carried out with rather coarse anatomical resolution, e.g., combining signal across the entire precentral gyrus, thus obliterating gradients between the hand and other motor areas that may have different roles in generating the low-beta and high-beta band activities. Here, we address these issues by applying a TDE-HMM based approach explicitly separating low- and high-beta activity and using finer anatomical graining, to capture functionally and anatomically distinct beta sub-bands.

Earlier studies on the reproducibility of the sensorimotor beta signal and beta events within individuals have treated beta as one homogeneous band (Illman et al., 2022; Niemelä et al., 2025; Pauls et al., 2024; Piitulainen et al., 2018). Moreover, the similarity of beta events across different measurement sites has not been examined. Larger datasets are increasingly being used in MEG data analyses (Cam-CAN et al., 2014; Power & Bardouille, 2021) and are necessary for, e.g., approaches that use machine learning or normative modelling (Bozek et al., 2023; Itälinna et al., 2023). Collecting large datasets is time-consuming and expensive, so pooling of data across measurement sites would be beneficial. Still, mixed-site cohorts continue to be rare, and site-dependent differences are frequently observed in data obtained from different neuroimaging modalities (Davatzikos, 2019). Previous MEG studies have suggested that pooling data across sites is feasible, as the inter-subject variability typically exceeds site variability (Boon et al., 2021; Ou et al., 2007) and measures such as source localization accuracy (Ashrafulla et al., 2011; Weisend et al., 2007) and source-based results (Hunt et al., 2019) are usually not affected by site. However, site variability likely depends on the measure and method of interest. In particular, the site variability of beta-related features across sites remains unknown and should be assessed before datasets can be combined, e.g., for normative modelling purposes.

We here used a TDE-HMM based approach to detect beta events from source-level MEG resting state data and examined its within-subject test-retest replicability across sessions, and its transferability between two measurement sites. Furthermore, we assessed the approach’s applicability to additional task data. To capture distinct activities within the beta band, we restricted the analysis to the beta band (13-30 Hz) and focused on the oscillatory characteristics during state detection (Korkealaakso et al., 2025). We compared the obtained separation of low- (< 20 Hz) and high- (> 20 Hz) beta states to the results obtained with an amplitude-based beta detection approach. We demonstrate that low- and high-beta state events have distinct occurrence probabilities and anatomical distribution at the source level which are reproducible across two sites and cohorts. Moreover, we demonstrate that the resting state model is transferable to a passive movement task, providing evidence of distinct modulation of high- and low-beta events which is not detectable using a solely amplitude-based approach. Furthermore, our results obtained from the same subjects across two different measurement days show that these measures have good test-retest reliability. In addition, the HMM-based approach yields comparable results when applied to resting MEG data recorded from two different MEG sites, showing that the approach is transferrable between sites.

## 2. Methods

### 2.1 Subjects

MEG data from a total of 84 healthy subjects were measured while they rested with their eyes open (restEO). 42 subjects (Aaltonen et al., 2023; Niemelä et al., 2025) were measured at site 1 (BioMag Laboratory), and 42 subjects (Nurmi et al., 2025) were measured at site 2 (Aalto University; subject characteristics summarized in Table 1). Additionally, 22 of the subjects measured at site 1 participated in a passive finger movement task (slowCKC), in which their left index finger was passively moved with a pneumatically driven device attached to the finger with an ISI of 3.5 seconds. After data preprocessing and artifact rejection, on average 86 trials per subject remained for analysis (median: 85.5 trials, range: 60-91 trials). Ethics statements were obtained from the HUS Regional Committee on Medical Research Ethics and the Aalto University ethics committee, and all participants gave their written informed consent before participating.

**Table 1.**
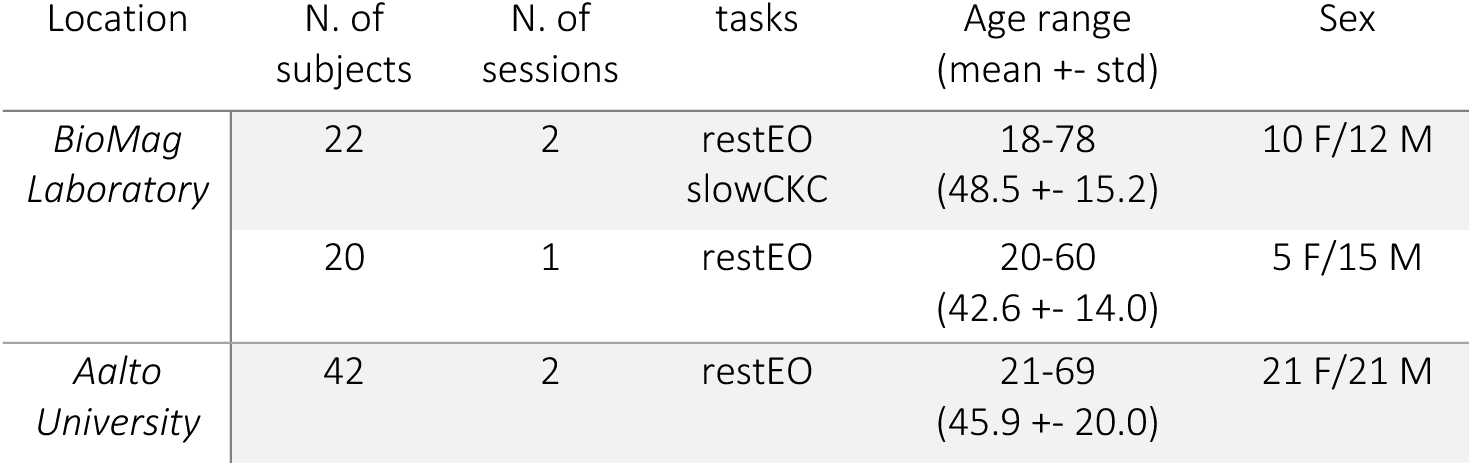
Summary of subjects included in the study.

| Location | N. of subjects | N. of sessions | tasks | Age range (mean +- std) | Sex |
| --- | --- | --- | --- | --- | --- |
| *BioMag Laboratory* | 22 | 2 | restEO<br>slowCKC | 18-78<br>(48.5 +- 15.2) | 10 F/12 M |
|  | 20 | 1 | restEO | 20-60<br>(42.6 +- 14.0) | 5 F/15 M |
| *Aalto University* | 42 | 2 | restEO | 21-69<br>(45.9 +- 20.0) | 21 F/21 M |

### 2.2 MEG recordings

The data were measured at the BioMag Laboratory at Helsinki University Hospital (site 1) and at the MEG Core at Aalto university (site 2). Both sites have a magnetically shielded room (site 1: ETS-Lindgren, Eura, Finland, site 2: Imedco AG, Hägendorf, Switzerland) with comparable 306-channel neuromagnetometers (site 1: Elektra Neuromag TRIUX, MEGIN Ltd, Helsinki, Finland, site 2: Elektra Neuromag Vectorview, MEGIN Ltd, Helsinki, Finland) consisting of 204 planar gradiometers and 102 magnetometers. To monitor head location during the recording, five head position indicator (HPI) coils were attached to the scalp and their locations were digitised with a 3-D digitiser pen (Fastrak, Polhemus, US). The 3-D digitiser was additionally used to digitise the subject’s head shape and three anatomical landmarks (the nasion and the right and left preauricular points). Spontaneous cortical activity was recorded with a 1-kHz sampling rate, together with continuous head position monitoring (cHPI). During the recordings, the data was band-pass filtered at 0.03-330 Hz. An empty room recording was available from the same day.

### 2.3 MEG data pre-processing and source reconstruction

For suppressing external artefacts, MEG data were pre-processed using the temporally extended signal space separation method (tSSS) (Taulu & Kajola, 2005), implemented in the MaxFilter software (MEGIN Oy, Helsinki, Finland, version 2.2.15). Subject’s head movements were corrected based on the cHPI recordings. Each subject’s head position was kept in individual space to avoid additional noise in the source reconstruction.

Further signal processing was done using MNE-Python version 1.7 (Gramfort, 2013; Larson et al., 2023). Artefacts arising from heart and eye movements were removed using the FastICA algorithm (Hyvarinen, 1999) implemented in MNE-Python based on visual inspection of the component time series and component topography. Artifacts originating from muscle activity remaining after ICA were removed by annotating the contaminated segments. The data was then filtered using a one-pass, zero-phase, non-causal band-pass FIR (finite impulse response) filter with lower and upper passband edges at 13 and 30 Hz. Finally, the data was downsampled to 200 Hz.

The bandpass-filtered and artefact-cleaned MEG data was then projected from the sensor space to the source space using FreeSurfer’s fsaverage MRI template as the structural space (Fischl, 2012). The fsaverage MRIs were individually scaled based on the digitised head shape points and the MEG-MRI transformation was carried out using the scaled fsaverage MRI. The boundary element model (BEM) for the forward model was created using the watershed algorithm (Ségonne et al., 2004) in MNE-Python. The forward solution was calculated using 5120 source points with fixed orientation perpendicular to the cortex (Hämäläinen et al., 1993). Inverse modelling was done using minimum norm estimate (MNE, signal-to-noise ratio **1.0**, loose **= 0**, depth **= 0.8**). Since the data was a continuous time series, the noise covariance was calculated using empty room recordings acquired on the same day, also applying tSSS and filtering (13-30 Hz) to the empty room data.

The dimensionality of the n=5120 source reconstructed MEG signals was reduced to n=450 parcels using the aparc_sub parcellation, a subdivided version of the aparc parcellation (Khan et al., 2018). Dimensionality reduction was carried out using pca_flip mode in MNE-Python, which best preserved the original time series’ oscillatory activity characteristics. The approach computes the first principal component across the sources within a label and applies a sign correction to avoid 180° phase changes.

### 2.4 Hidden Markov model for characterizing beta events

Applying the Hidden Markov Model (HMM) framework to MEG data allows decomposition of the MEG time series data into a sequence of state activations (Vidaurre et al., 2016a; Vidaurre, Hunt, et al., 2018), where each state has a probability distribution estimated from the data. The inference estimates the probability of each sample (i.e. time point) to be generated by each state’s estimate, given past samples. Here, we used the Time-Delay Embedded (TDE) model (Vidaurre, Hunt, et al., 2018), which instead of only focusing on one observation ***Y*_*t*_** at time point ***t***, models the joint distribution of an embedding of points from ***t* − *L*** to ***t* + *L*** (here with a 85-ms time window) using a (zero-mean) Gaussian distribution. The key state parameter is therefore a covariance matrix across time and space (i.e. an autocovariance) which characterizes the spectral properties of the signal within the chosen time window. Specifically, the observations (MEG data points) become ***Y*_*t*_ = (*y*_*t*-*L*_, …, *y*, …, *y*_*t*+*L*_)** and the TDE observation model becomes

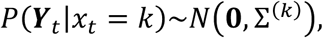

i.e., each state ***k*** is represented by an autocovariance matrix **Σ^(*k*)^**. We determined the appropriate number of states ***K*** by testing different models varying ***K*** from 2 to 6, with the aim of optimising the separation between states with different spectral content. Based on previous literature, our prior was that the model should be able to distinguish two oscillatory states within the beta band (13-30 Hz). Oscillatory activity was best represented by ***K* = 4** states, and increasing ***K* > 4** did not result in additional, better frequency separation in the beta band (see **Supplementary Figure 1**). After initial testing, we set the number of states to be ***K* = 4** and the lag parameter to be ***L* = 8** with step size being 1. The estimation of the posterior distribution parameters is inferred using variational Bayes and stochastic inference (Vidaurre, Abeysuriya, et al., 2018; Vidaurre et al., 2016b).

We first fit the HMM uniformly for all subjects and parcels, i.e., at the group-level. All ***N*** subjects with ***P*** parcel-level time courses of ***T*** timepoints were concatenated, yielding a dimensionality of **ℝ^*NPT*×1^** for the group-level model. The group-level model’s states are common for each subject, meaning that the states’ probability distribution parameters are obtained uniformly for all subjects and parcels. This assumes that similar activity falls into the same state in all subjects, ensuring that the states are comparable between subjects. In a second step, we re-estimated the group-level HMM separately for each subject and parcel to obtain subject- and parcel-specific state parameter estimates. This process, referred to as dual estimation, ensures the comparability of states across subjects and parcels (Vidaurre et al., 2017). We used the Matlab version of the HMM-MAR toolbox for data analysis (Vidaurre et al., 2016b); a Python implementation has also recently been made available (Vidaurre et al., 2025).

Given MEG data as input, the HMM process results into state probability time courses, in which each state has a probability to be active at each time point. Based on these state time courses and the estimated state transition probabilities, a binary time course of the active states can be derived via the Viterbi algorithm (i.e., Viterbi path). We used the state time courses and the Viterbi path to derive additional features from the time series data.

### 2.5 Beta characteristics extraction

The outputs of the individually estimated HMMs are (1) the probability time courses and (2) the Viterbi path. The former gives the probability of each state to underlie the observed data at each timepoint, whereas the latter is a binary time course of the most likely sequence of state activations. Using these model outputs and MEG parcel time series, we calculated several state characteristics, namely fractional occupancy (FO), state lifetime (SLT), event rate, and event amplitude (EA). The fractional occupancy describes the proportion of each state being active of the total recording time, calculated by dividing the time spent in a state by the total duration of the time series. The state lifetime refers to the event duration in milliseconds (per state visit) and the rate to the number of events of one state occurring per second. SLT was calculated by taking the mean of all state visits longer than 50 ms, separately for each state. The rate was derived by dividing the total number of state visits per state by the total duration of the resting data, giving an estimate of events per second. EA describes the signal’s amplitude during a state visit. To calculate EA, the original 13-30 Hz band-filtered parcel MEG time series was decomposed with a set of complex Morlet wavelets within the frequency range with 1 Hz resolution and ***n*_*cycles* = *frequency*/2**. Then, the mean of the absolute value of the signal’s time-frequency representation was taken over the 13-30 Hz range, resulting in a beta range amplitude envelope. Finally, the mean of all timepoints within a state visit exceeding the 75^th^ percentile of the values was calculated for each individual state visit. States’ PSDs were calculated using the multitaper method with 7 tapers and time-bandwidth product of 4 (Vidaurre et al., 2016b).

### 2.6 Conventional beta amplitude envelope analysis

In addition to the HMM based beta characteristics, we also calculated the conventional beta amplitude envelope-based characteristics for comparison. The amplitude modulation was calculated for each cortical area (450 parcels) independently. This was done using the same beta band-pass filtered data (13-30 Hz). The amplitude envelope was calculated similarly to the HMM-based EA characteristic, such that the band-filtered parcel MEG time series was decomposed with a set of complex Morlet wavelets within the frequency range with 1 Hz resolution and ***n*_*cycles* = *frequency*/2**. Then, the absolute value of the signal’s time-frequency representation was estimated separately for the low- and high-beta frequency ranges (13-20 Hz and 20-30 Hz), resulting in a high- and a low-beta band amplitude envelope. The amplitude of the beta events, defined as periods exceeding the 75^th^ percentile amplitude threshold, was averaged per band. High- and low-beta amplitude envelopes were used to calculate the induced responses without prior thresholding (see section 2.7). The PSD for the whole beta band (13-30 Hz) was calculated for the original parcel-level time series similarly to the HMM state PSDs using the multitaper method with 7 tapers and time-bandwidth product of 4.

### 2.7 Task- related state probability and amplitude modulation

Next, we analysed state characteristic modulation during the left finger passive movement task. For the HMM-based states, we calculated task-related state probabilities by averaging the state probability time courses across trial epochs (-0.5 to 3 s relative to the onset of the passive movement). The task-related state probabilities were individually normalized by the average state probability during rest, and the baseline interval (-0.5-0 s) was set to zero.

Next, the mean across all subjects was calculated to obtain a group average state probability curve. A two-sided non-parametric cluster-level paired t-test for temporal data, as implemented in MNE-Python, was used to test if the task-related state probability curves differed from baseline interval (-0.5-0 s) (number of permutations 1024, degrees of freedom ***n*_*subjects* − 1**, cluster forming threshold ***p* < 0.001**, significance threshold ***p* < 0.001**). Here, the multiple comparison problem is addressed with a cluster-level permutation across time.

To assess conventional beta amplitude envelope modulation during the task (in comparison with the HMM-based approach), we calculated the signals’ amplitude envelopes for low-beta (13-20 Hz) and high-beta bands (20-30 Hz), as described for resting data in section 2.6. The resulting amplitude envelopes were segmented into epochs (-0.5 to 3 s relative to the onset of the passive movement) and averaged to obtain the induced responses. Similarly to the task-related state probabilities, the induced responses were then individually normalized by the average amplitude envelope during rest, and the baseline interval (-0.5-0 s) was set to zero. The group average was then taken as a mean over the subjects. A two-sided non-parametric cluster-level paired t-test for temporal data, as implemented in MNE-Python, was used to test if the task-related state probability curves differed from baseline interval (-0.5-0 s) (number of permutations 1024, degrees of freedom ***n*_*subjects* − 1**, cluster forming threshold ***p* < 0.001**, significance threshold ***p* < 0.001**). Here, the multiple comparison problem was addressed with a cluster-level permutation across time.

We also investigated the spatial differences between the states’ probability time courses and frequency bands amplitude modulation, described in detail in **Supplementary methods**.

### 2.8 Parameter stability and model transferability

To assess the test-retest reliability of the extracted characteristics (FO, SLT, rate and EA), the same group model was applied to data from two separate MEG sessions obtained 3 months apart in 22 subjects at site 1. The test-retest stability was estimated with intraclass correlation (ICC), as implemented in the Python Pingouin package (Vallat, 2018). We used an ICC variant with two-way mixed effects with consistency *ICC*(3,1), which becomes

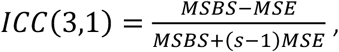

where *MSBS* and *MSE* are the mean square between subjects and mean square error, respectively, and *s* is the number of sessions. The obtained ICC values were classified as poor **< 0.4**, fair **0.4 − 0.59**, good **0.6 − 0.74** or excellent consistency **0.75 − 1.00** (Cicchetti, 1994).

Group model transferability between the two sites was addressed by applying the group model derived from site 1 data to the data measured at site 2 (different MEG device, measurement environment, and subject cohort). The same characteristics (mean FO, mean SLT, rate and mean EA) were then extracted for the site 2 data. We assessed the difference between the extracted resting state characteristics separately for each parcel with a parametric two-sided t-test with *p* < 0.05.

### 2.9 Code and model availability

The pretrained model and research code are available at https://github.com/BioMag/beta_event_characterisation.

## 3. Results

### 3.1 The HMM-based pipeline decomposes two distinct beta brain states and provides additional information compared to an amplitude-envelope based analysis

The HMM process with uniform fit for all subjects and parcels, i.e., at the group-level, resulted in a 4-state group model (spectral behaviour shown in Figure 1A). Model stability was confirmed by generating several model permutations using different subsets of subjects, as well as generating a model using data from site 2 (different site and cohort, see Supplementary Figure 4). The model identified two states with periodic oscillatory activity in the beta band (states S1 and S3), and two additional states (states S2 and S4). Of the two beta-band oscillatory states, one has a PSD frequency peak below 20 Hz (low-beta, S3) and the other one has a frequency peak above 20 Hz (high-beta, S1). The HMM-based low- and high-beta PSDs match the low-beta and high-beta periodic activity peaks in the original PSD from the parcel’s timeseries (see **Figure 1**B for an example in a single parcel). The other states’ (S2 and S4) PSD patterns do not show oscillatory activity in the beta range and likely represent activity remaining outside beta band after the band pass filtering at 13-30 Hz (Korkealaakso et al., 2025). The PSD of S4 increases towards the lower band boundary of the PSD, compatible with activity in the sub-beta band (below 13 Hz) and thus referred to here as sub-beta state (see also Korkealaakso et al., 2025). S2 has no periodic activity in the beta-range: Based on our previous findings (Korkealaakso et al., 2025), we postulate that state S2 corresponds to gamma activity (henceforth referred to as gamma state). We here focus on the two beta states. The properties of the states and their distinct modulation during an active motor task are shown more comprehensively in Korkealaakso et al., 2025.

**Figure 1.**
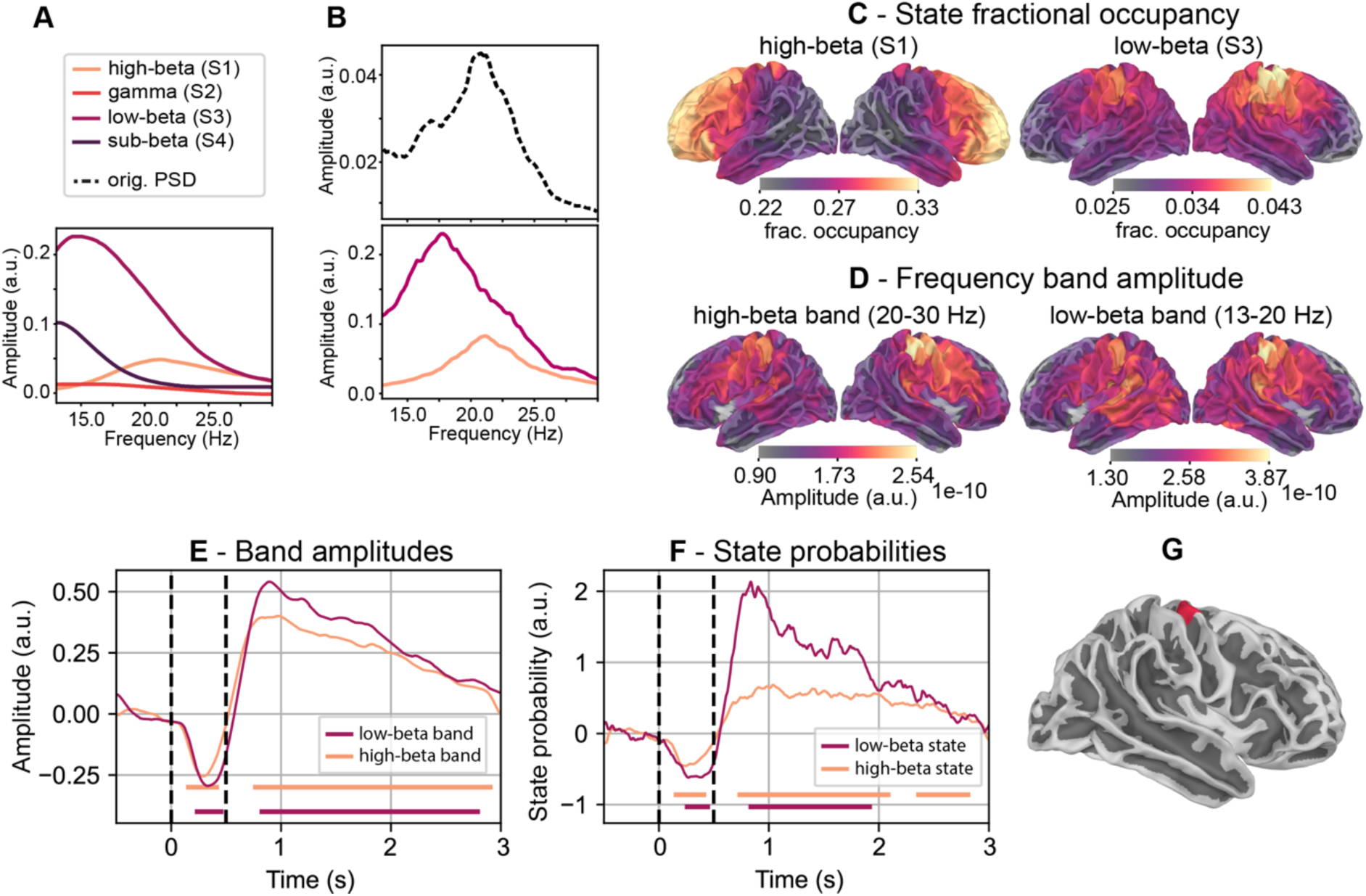
**A.** PSD of the states from the group model and **B.** an example of one individual’s PSDs at the right-hemispheric pre­central parcel (location shown in G). **C.** Resting group-level fractional occupancies of the low- and high-beta states showing distinct distributions across the brain. **D.** Resting mean amplitude envelope for the low-beta (13-20 Hz) and high-beta (20-30 Hz) bands. Here, only the magnitude of the two beta band amplitudes differs, while the anatomical distribution is very similar. **E.** Induced activity during the passive finger movement task in the right hemisphere pre-central parcel (shown in **G**) for the high- (20-30 Hz, orange) and low-beta (13-20 Hz, violet) bands. The induced response is derived relative to the resting state amplitude, and the baseline interval is set to zero. Periods differing significantly from baseline interval (-0.5-0 s) are indicated by same-coloured bars below the curves, *p* < 0.001. **F.** State probability modulation during the passive finger movement task in the right hemisphere pre-central parcel (shown in **G**) for the high- and low-beta states. The state probability is derived relative to the resting state probability, and the baseline interval is set to zero. Periods differing significantly from baseline interval (-0.5-0 s) are indicated by same-coloured bars below the curves, *p* < 0.001. **G.** Anatomical location of the selected right hemisphere parcel in the pre-central area.

The fractional occupancy (FO) at rest, i.e. the proportion of the total time series occupied by each state, shows a distinct anatomical distribution for the two beta states (**Figure 1**C, see also **Supplementary Figure 2** for the other states’ anatomical distribution). The low-beta state FO maximum is centred around the sensorimotor cortex, whereas the high-beta state occurs most frequently in the frontal areas. The two beta states’ FOs differ greatly: In areas where high-beta is most prevalent, it occurs about 30% of the time, while the corresponding occurrence for low-beta is only 5% in its own most prevalent areas. Low-beta events are therefore very rare compared to high-beta events.

The HMM-based analysis and amplitude-envelope based analysis provide complementary results (see Figure 1C and Figure 1D): While the HMM-based analysis gives the proportion of time of a certain type of oscillatory activity occurring, amplitude-envelope based results reflect changes in the signal amplitude in a specific oscillatory band. State FO for the two beta states is very distinct anatomically (Figure 1C), whereas the anatomical distributions of the two beta frequency bands obtained from the amplitude-envelope based approach look almost identical (Figure 1D), resembling the event amplitude distribution obtained from the HMM-based approach (Supplementary Figure 2). The maximum of amplitude envelope-based beta-band resting-state activity localizes to areas surrounding the central sulcus for both low-beta (<20Hz) and high-beta (>20Hz) bands (Figure 1D), with lower signal magnitude for the high-beta band. Interestingly, the anatomical distribution of event amplitudes obtained from the HMM-based approach is almost identical for all four states (Supplementary Figure 2B). These findings demonstrate that amplitude-envelope based approaches bias towards areas with higher amplitudes. The HMM-based approach, detecting the probability of different oscillatory frequency patterns, thus adds information by also revealing lower-amplitude oscillatory activity.

Next, we assessed transferability of the HMM resting state-based group model to event-related activity recorded during a passive finger movement task and compared the results with conventional induced activity of the event-related data based on an amplitude envelope approach. Figure 1E shows the amplitude envelope-based induced responses, demonstrating a typical beta suppression-rebound pattern which looks identical for both beta frequency bands, differing only in magnitude. In contrast, Figure 1F shows the state probability modulation of the low- and high-beta states’ during the same task in the same right hemisphere pre-central parcel, also revealing modulation in a suppression-rebound manner, but with marked differences between the states: During the rebound phase (0.5-3.0 s), low-beta probability increases almost twofold compared to rest, demonstrating a pronounced low-beta rebound after passive movement. In comparison, high-beta state modulation is much smaller during the rebound. Thus, although the HMM state model used here was initialized using resting data in a data-driven manner (and therefore task-agnostic), the states captured brain activity relevant to motor function and behaviour. Anatomical patterns of the HMM-based and amplitude-envelope based analyses are shown in the supplementary materials (see **Supplementary Figure 3**), demonstrating that low-beta probability is modulated over a wider cortical area than high-beta, which is not visible in the amplitude envelope-based analysis.

### 3.2 State characteristics show good to excellent test-retest stability

Next, we calculated within subject test-retest stability for the state characteristics across two measurement sessions. The results are shown in **Figure 2**A for the high-beta state and in **Figure 2**B for the low-beta state. For high-beta FO, SLT and EA, most of the parcels have ICC values **> 0.75**, corresponding to excellent consistency (FO: 448/450 areas, SLT: 415/450 areas, EA: 440/450 areas; Figure 2A). Only event rate is slightly less stable in the frontal areas, but the sensorimotor areas have excellent consistency (385/450 areas with ICC > 0.75). Low-beta state characteristics are less stable, possibly due to low-beta’s low FO (about 5%): the low-beta state is therefore present for only 10-15 seconds during a 3-minute resting state measurement. For the low-beta state, only the event amplitude shows excellent consistency throughout the cortex (427/450 areas; Figure 2B). The SLT and rate have mostly good to fair ICC, while the ICC of the FO is excellent in some areas, e.g., in the right motor cortex (ICC fair or better, FO: 398/450 areas, SLT: 322/450 and rate: 246/450 areas with good or fair consistency).

**Figure 2.**
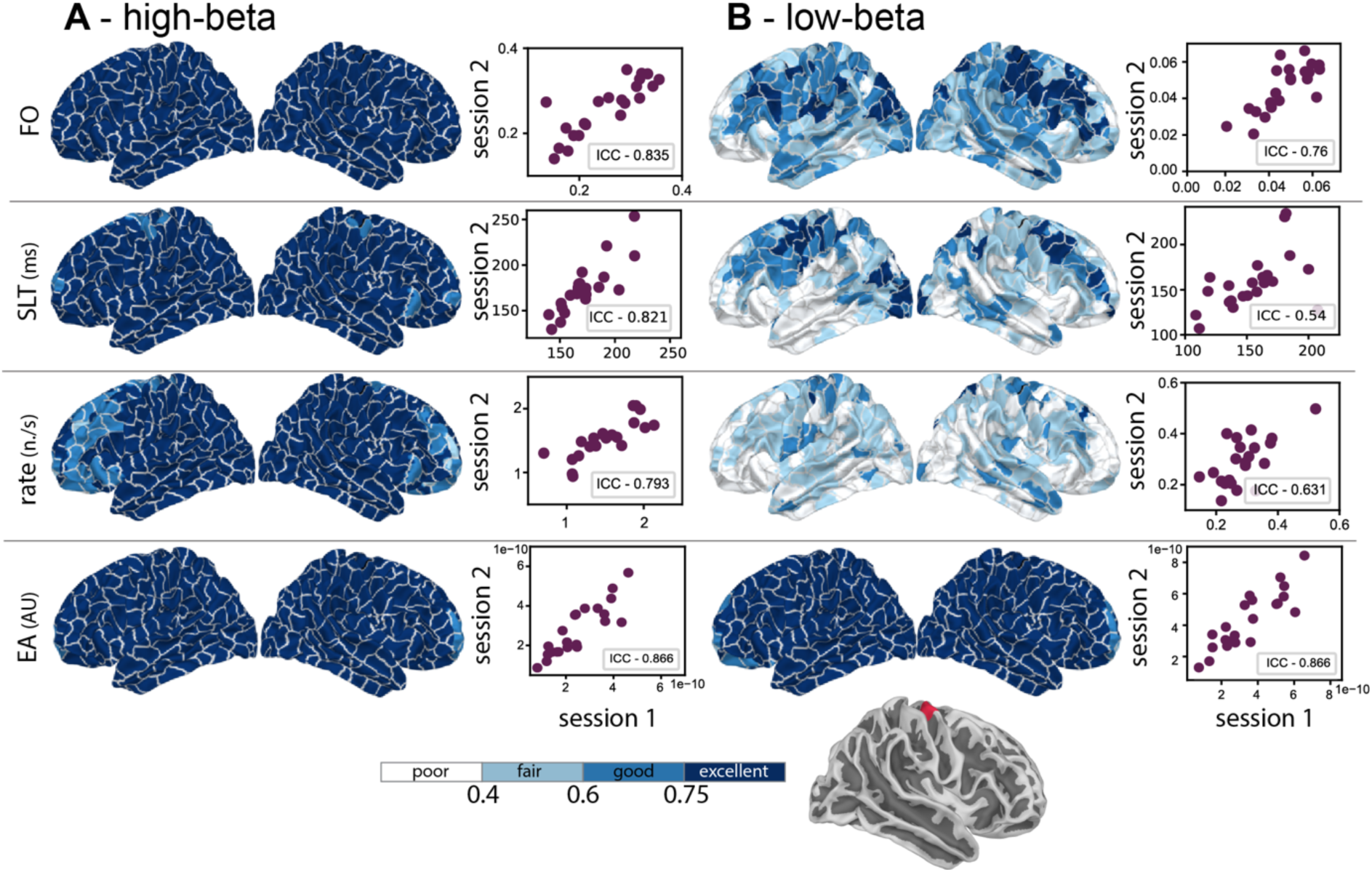
Test-retest stability of state characteristics. Stability of the high-beta (**A**) and low-beta (**B**) states across sessions. Intra-class correlation (ICC) was used as the stability measure; the stability was calculated for fractional occupancy (FO), state lifetime (SLT), event rate and event amplitude (EA) from two separate resting state measurement sessions 3 months apart. The brain representation (left) maps the ICC values from all parcels (ICC value legend at the bottom), whereas the right-panel scatter plots show the distribution of individual values of each measure across subject in a single parcel (right hemisphere pre-central gyrus parcel visualized at the bottom).

### 3.3 The group model is reproducible and transferable between two measurement sites and cohorts

To assess whether the HMM group model can be transferred across sites, we applied the group model trained on site 1 data to data from site 2. **Figure 3** shows the results for the high-beta and low-beta states. The data features show highly similar anatomical distributions across the two sites, hence reproducing our results in a second, independent subject cohort and site. Quantitatively, there were hardly any statistically significant differences between the FO, SLT and event rate values between the sites, i.e., most of the parcels’ characteristics did not differ (two-sided t-test with ***p* < 0.05**); the few parcels with significant differences are indicated in orange in the middle two columns in Figure 3. However, while the anatomical distribution of EA was qualitatively very similar for both beta states and both sites, quantitatively it differed in between the two sites in many brain areas (see Figure 3 bottom row). Site differences were statistically significant in the frontal, occipital and temporal areas. These areas have lower signal amplitude in the beta range, which can make them susceptible to signal-to-noise ratio issues. In all measures not relying on raw data, the site transferability was good, suggesting an HMM-based analysis approach may be advantageous over amplitude envelope-based estimation when comparing or combining subject cohorts from different sites.

**Figure 3.**
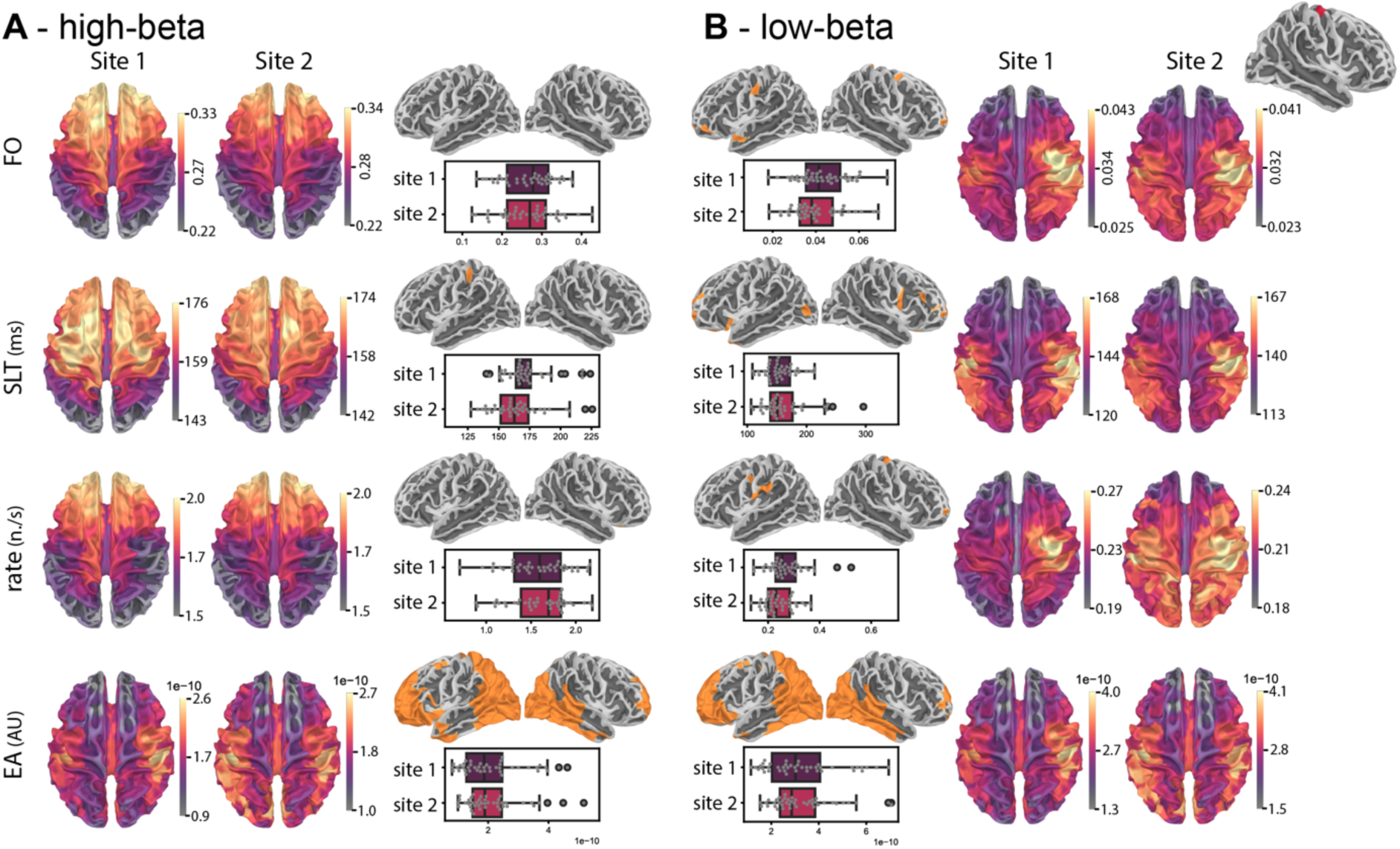
Group model applied to data measured at two sites from two independent cohorts. Characteristics (FO, SLT, rate and event amplitude) shown in consecutive rows for site 1 and site 2 for the high-beta (**A**) and the low-beta (**B**) states. The anatomical renderings (the left-most and right-most panels) show group maps of each parameter’s distribution for the two different sites. The two middle panels show parcels (orange) with statistically significant differences between the sites (t-test p<0.05). Boxplots show how individual subjects’ values are distributed within one parcel over the two data sets (right hemisphere pre-central parcel visualized in the brain inset at the top right).

## Discussion

Our results demonstrate that the proposed HMM-based approach can decompose beta band activity in MEG data into low- and high-beta states in a data-driven manner. These findings were reproduced across two independent subject cohorts, highlighting the robustness of the approach. Moreover, the model generalizes to task data and provides complementary information to conventional amplitude envelope-based approach, underscoring the advantages of characterizing activity via frequency-driven state probabilities. Several model-derived features exhibit good to excellent within-subject reproducibility across measurement sessions. As the trained model produces comparable results across different measurement sites, it allows, e.g., pooling of data to increase cohort sizes. Finally, the approach eliminates the need for manual, observer-dependent decisions, thus enabling automated whole-brain analyses at the source level.

Frequency-driven HMM applications to MEG data (Vidaurre, Hunt, et al., 2018) have previously been used to characterize whole-band beta events both at the single-channel (Seedat et al. 2020) and network level (Kohl et al., 2025). Given the evidence of sub-bands within the wide beta band with probably different functional roles (Barone & Rossiter, 2021; Cao et al., 2024, Nougaret et al. 2024), we aimed to decompose activity to sub-beta bands at the single-parcel level. The resulting two sub-bands had distinct and reproducible spectral characteristics and anatomical distribution across two independent subject cohorts.

Furthermore, the two beta states also showed distinct state-specific modulation during task. The high-beta state was present most prominently in frontal areas, whereas low-beta occurred most frequently in motor areas, which is in line with previous findings of a posterior-to-anterior gradient of increasing beta frequencies (Capilla et al., 2022).

Apart from the obvious benefits of automated analysis in terms of speed and elimination of observer bias, removing the relative weighting of signal amplitude can bring out new features in the data. We found that MEG signal amplitude was anatomically similarly distributed across all states and frequency bands with a maximum around the central gyrus, while fractional occupancy and its distribution were very distinct for the high- and low-beta states. Signal amplitudes are usually higher for lower frequency oscillations, thus making detection of, e.g., gamma (30-90 Hz) activity more challenging compared to lower frequency activity.

Thus, focusing on frequency characteristics, as proposed here, may improve feature detection for measures obtained from raw data, and for analyses targeting higher frequency oscillations, especially outside regions surrounding the central gyrus.

Test-retest stability is critical, e.g., for potential diagnostic approaches. Beta modulation during motor tasks (Illman et al., 2022; Niemelä et al., 2025), its resting-state characteristics (Martín-Buro et al., 2016; Pauls et al., 2024), and the modulation of beta-band connectivity during language processing (Ala-Salomäki et al., 2025) have been shown to be stable. In agreement with previous findings, we here show that beta events’ amplitude is very stable for both high- and low-beta states at rest, indicating that amplitude has a generally stable nature across the entire brain at the beta band. High-beta state characteristics appeared very stable across measurements, while low-beta showed more modest test-retest reliability. This difference may partly reflect the low occupancy of the low-beta state, which could lead to relative undersampling during the short (3-min) recordings used here, thereby reducing the reliability of the estimates. In addition, beta events have been shown to propagate across the cortex at rest (Zich et al., 2023), potentially causing inherent spatial and temporal variability. Such intrinsic dynamics may also limit the stability of low-beta measurements across sessions. Future studies using longer resting- state measurements are needed to clarify this issue.

In many settings, it would be beneficial to pool MEG data from multiple locations, e.g., when studying rare patient populations, or to enable, e.g., machine learning and normative modelling approaches (Bozek et al., 2023; Itälinna et al., 2023; Vaghari et al., 2022), which require large amounts of data. Here, we tested the transferability of the model to identify characteristics from data collected at two different MEG sites with different cohorts. We show very good site and cohort transferability for all state characteristics except for event amplitude. This may reflect differences in noise levels of the MEG devices or shielded rooms, or different measurement practices between locations. The observed site-specific differences in amplitude also underline the potential benefits of approaches and measures that are less affected by signal amplitude when combining or comparing data across multiple sites, such as the one we here propose.

The proposed HMM-based approach relies on modelling choices, such as the number of states and the use of narrow-band filtering, which need to be considered when interpreting the results. Based on biological priors, we here chose a four-state model for maximizing the separation of two distinct beta-range states. The subsequent analysis thus constrains every data point to one of the four states. While the discretization is intentional, it applies a specific classification scheme for the data. As the optimal classification of the signal is not known a priori, our model should be viewed as a solution that shows meaningful dynamics from the data with respect to beta band, rather than the only solution. The chosen model produced corresponding anatomical maps for two independent subject cohorts without incorporating prior anatomical information. We restricted the signal to the beta band to ensure that classification specifically reflects activity in this range. However, this choice may be limiting in certain contexts, such as patient cohorts with spectral slowing or atypical spectral profiles. In such cases, it will be important to examine how the model classifies altered (and known pathological) activity to avoid potential misclassification and misinterpretation.

In conclusion, the proposed frequency-based HMM approach decomposes beta range into two distinct beta bands in a time-resolved manner and reveals new features in the data, thus extending findings from amplitude envelope-based analysis approaches. The approach is transferable to task data and shows task-related modulation differences between high- and low beta which are not detectable with amplitude envelope-based methods. In addition, the approach is stable across different measurement sites and cohorts, facilitating, e.g., joint analysis of multi-site data sets.

## Funding and acknowledgements

CA was funded by a Carlsberg Foundation Visiting Postdoctoral Fellowship at the University of Oxford (CF23-1716). DV was supported by a Novo Nordisk Foundation Emerging Investigator Fellowship (NNF19OC-0054895) and an ERC Starting Grant (ERC-StG-2019–850404). ML was supported by the Swedish Cultural Foundation in Finland, the Sohlberg Foundation, and the Finnish Cultural Foundation. HR received funding from the Research Council of Finland (Grant Nos. 355409 and 321460 and Flagship of Advanced Mathematics for Sensing Imaging and Modelling grant 359181) and the Sigrid Juselius Foundation. KAMP was in part supported by the Research Council of Finland (grant number 350242), the Sigrid Juselius Foundation, the Finnish Medical Foundation, and a government research grant (valtion tutkimusraha).

The authors thank all study participants for providing their time and Linda Niemelä, Jari Kainulainen, Juho Aaltonen, Hanna Kaltiainen and Pietari Nurmi for measuring the data sets used in the analyses.

## Code availability

The pretrained group model and research code are available at https://github.com/BioMag/beta_event_characterisation.

## Supplementary materials

### Supplementary methods

#### Spatial patterns of modulation of the task-related features

The spatial properties of the task-related state probability modulation (see **Supplementary Figure 3,** D-F) were studied by looking the difference during suppression versus rebound phases compared to resting activity. Activity during suppression phase was considered as mean probability over 0.1–0.4 s and rebound was derived as mean probability over 0.6-1.6 s of the task-related state probability (see section 2.7), resulting in state probability modulation during the task relative to rest.

In analogy to the HMM-based state probability modulation, the spatial properties of the induced responses were studied by calculating difference during suppression versus rebound phases (see Supplementary Figure 3, A-C) of the induced response (see section 2.7) for the same suppression and rebound time intervals, compared to rest, resulting in power modulation during the task compared to rest.

Differences between the states’ (HMM-based) and bands (amplitude envelope based) spatial suppression-rebound modulations were tested with two-sided parametric t-test and Bonferroni correction (t-test threshold ***p* < 0.001**, Bonferroni correction alpha level ***α* < 0.05**, see **Supplementary Figure 3** C and F).

## Supplementary figures

**Supplementary Figure 1.**
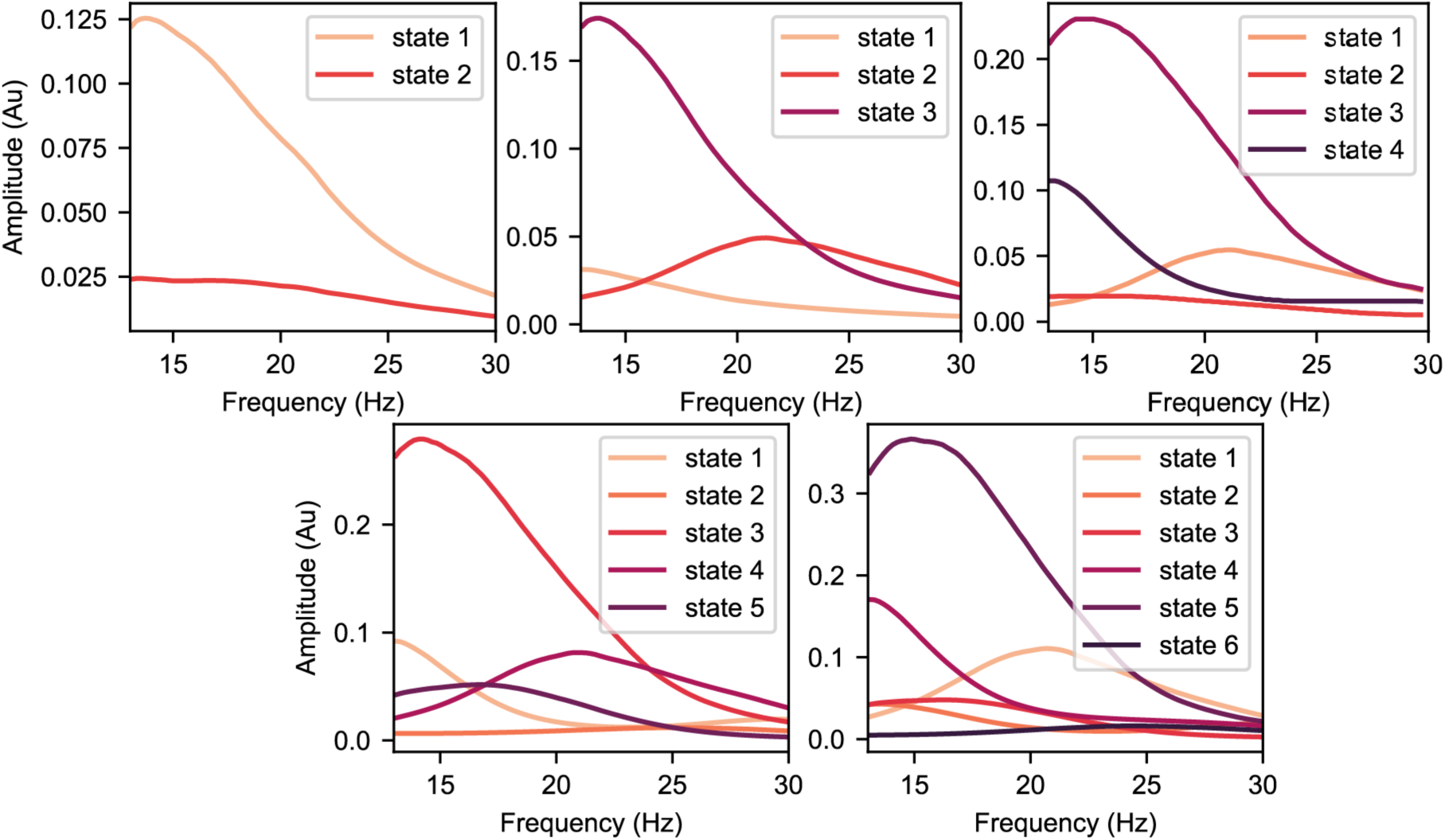
Power spectral densities for group model with *K* = 2, 3, 4, 5 and 6 states.

**Supplementary Figure 2.**
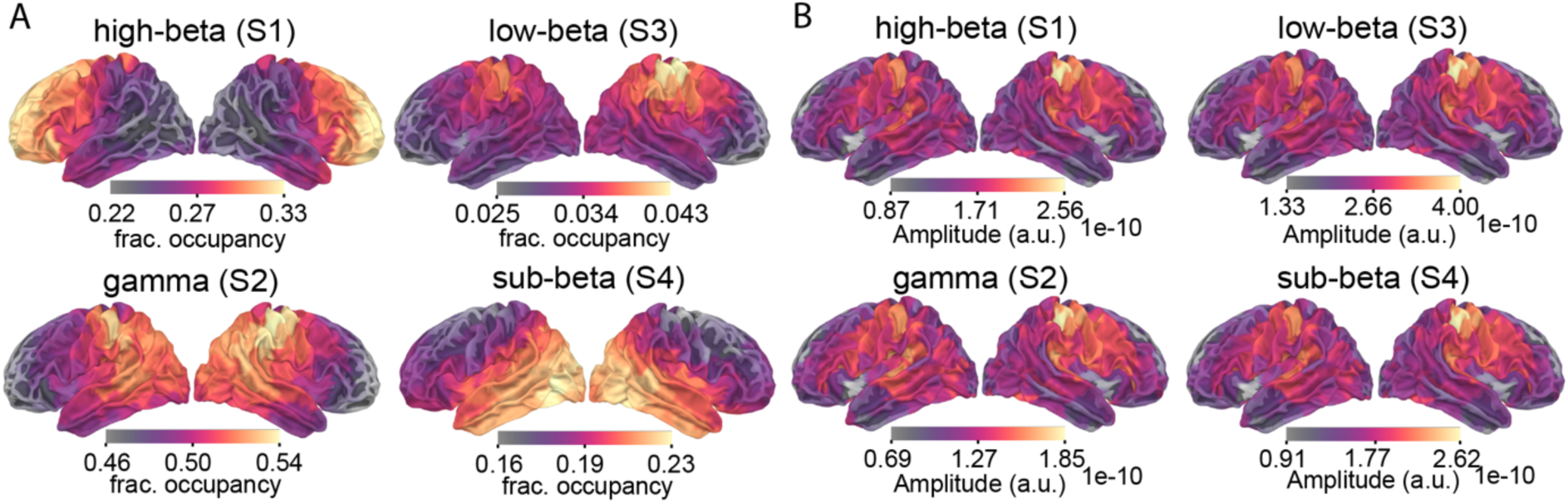
Anatomical distribution of **A.** fractional occupancy for the four states and **B.** their respective mean event amplitudes.

**Supplementary Figure 3.**
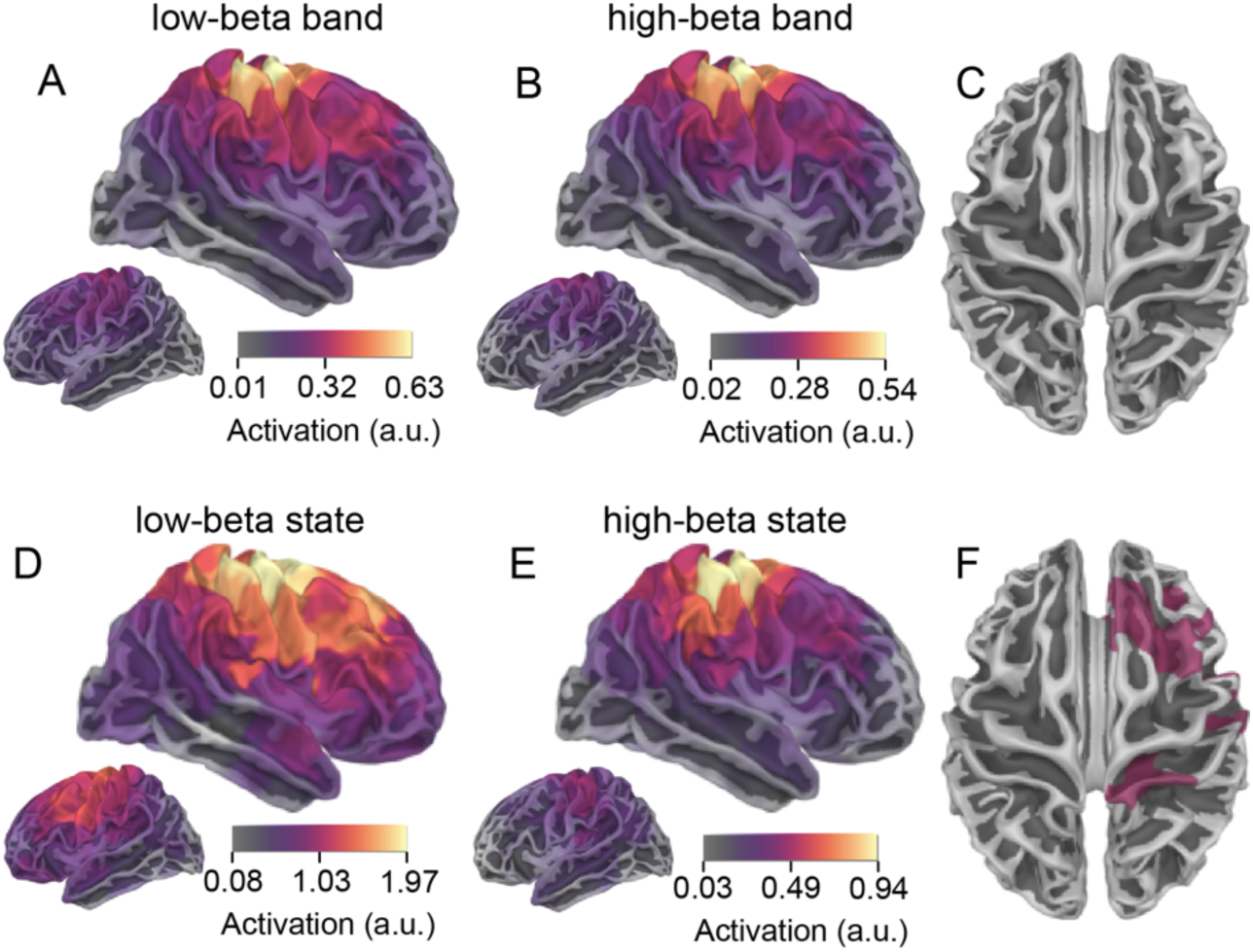
Anatomical distribution of HMM versus amplitude envelope derived task related features’ modulation. Anatomical areas with significant modulation of the low-beta (**A.**) and high-beta (**B.**) bands’ suppression-rebound difference in induced power (amplitude envelope approach) relative to rest (rebound: mean during 0.6-1.6s, suppression: mean during 0.1-0.4s). **C.** Anatomical areas with statistical differences between both beta bands’ right hemisphere’s modulations (significant modulation shown as coloured areas, two-sided t-test with ***p* < 0.001** and Bonferroni correction ***α* < 0.05**). Anatomical areas with significant state probability modulation of the low-beta (**D.**) and high-beta (**E.**) states’ suppression-rebound difference relative to rest. **F.** Anatomical areas with statistical differences between the states’ right hemisphere’s modulations (significant modulation shown as coloured areas, two-sided t-test with *p* < 0.001 and Bonferroni correction *α* < 0.05).

**Supplementary Figure 4.**
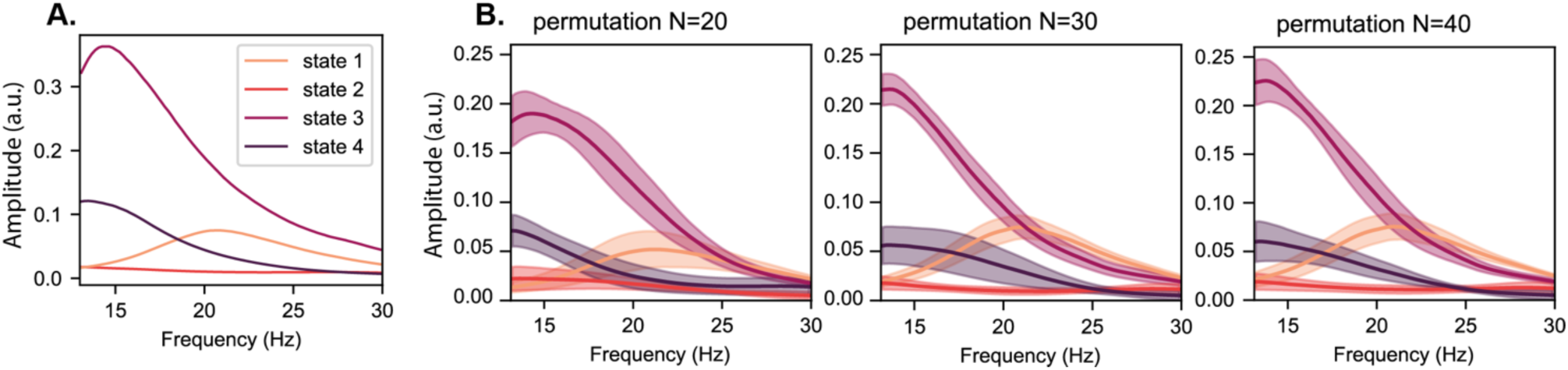
Convergence of the four-state model. **A.** Power spectral density for group model generated using site 2 data. **B.** Power spectral density average for model permutations (10 models) using different subsets of subjects (number of subjects N=10,20 and 30) from site 1. The shaded area shows one standard deviation from the mean.

## Notes

### Competing Interest Statement

The authors have declared no competing interest.

